# Sex-specific trajectories of gait adaptation following partial dopaminergic lesions in a rat model of parkinson’s disease

**DOI:** 10.1101/2025.11.10.687728

**Authors:** Diego Lievano Parra, Juan D Garavito, Greg Jensen, Valeria V González, Fernando P Cardenas

## Abstract

Parkinson’s disease (PD) is a neurodegenerative disorder in which gait disturbances are central to disability, yet their early manifestations remain poorly understood due to the difficulty of identifying prodromal PD in humans. Rodent models may help narrow this gap by approximating partial dopaminergic loss and enabling investigation of motor changes that precede overt symptoms. Here, we examined gait dynamics in Wistar rats following unilateral, low-dose 6-hydroxydopamine (6-OHDA) lesions in the substantia nigra pars compacta (SNc). Animals were assessed weekly for six weeks in the horizontal ladder test, where running velocity and footfall accuracy were quantified with markerless pose estimation using DeepLabCut (DLC). Lesioned animals exhibited greater tyrosine hydroxylase (TH)-reactivity asymmetry than controls, reflected behaviorally as persistent but gradually attenuating deficits in gait precision. Errors peaked at week three and remained above baseline through week six. Velocity was preserved across groups, but sex differences emerged. Control females accelerated more steeply over time, whereas lesioned females showed attenuated gains and greater variability, while males displayed more homogeneous velocity profiles across conditions. Together, these findings indicate that partial SNc lesions reveal sex-specific trajectories of motor adaptation, a feature relevant to modeling prodromal PD. Combined with DLC-based tracking, this framework offers a practical approach for detecting early behavioral markers and supporting the identification of preclinical motor features of PD.

## INTRODUCTION

Parkinson’s disease (PD) is a neurodegenerative disorder that affects approximately 2% of individuals over the age of 60 and is clinically characterized by resting tremors, rigidity, bradykinesia, and gait abnormalities^[1,2]^. While these motor symptoms are well described in advanced stages ^[3]^, the impact on motor behavior during early stages remain poorly understood. Identifying the onset of neurodegeneration is particularly difficult because pathophysiological changes may precede quantifiable motor abnormalities. Because progressive dopamine depletion is a defining pathological feature of PD, rodent models that replicate partial loss of dopaminergic neurons can offer unique opportunities to investigate motor alterations that precede overt clinical diagnosis.

Among these models, the 6-hydroxydopamine (6-OHDA) model has been widely used to mimic PD-like motor deficits in rodents, showing a clear dose-dependent relationship between dopamine depletion and motor impairment ^[4]^. Traditionally, high-dose of 6-OHDA have been used to produce extensive lesions, reliably reproducing the motor deficits that resemble advanced PD ^[5,6]^. Nevertheless, these approaches provide limited insight into the subtle and dynamic behavioral changes that accompany early stages of dopamine depletion.

To address this limitation, low-dose 6-OHDA protocols can induce moderate lesions that mimic mild, variable, and adaptive motor deficits, offering a closer approximation to early stages of PD ^[7]^. Furthermore, targeting the substantia nigra pars compacta (SNc) is advantageous, as it captures the earliest site of dopaminergic vulnerability in PD ^[8]^. Nevertheless, behavioral outcomes of moderate lesions can be variable, due to factors such as lesion site, dosage, assessment window, and subject characteristics ^[9]^. For instance, moderate dopaminergic depletion in the medial forebrain bundle (MFB) and dorsal striatum (DS) often leads to measurable gait impairments ^[10]^, whereas similar depletion in the SNc may produce subtler effects ^[11]^. This variability underscores the importance of refining SNc-targeted lesions, as they offer a promising framework for modeling motor patterns that resemble early PD.

In addition, most studies have focused on male rodents, leaving sex-specific adaptations largely unexplored, despite evidence that non-dopaminergic factors modulate dopaminergic plasticity, compensation, and motor outcomes ^[12]^. Consequently, distinct and progressive patterns of gait adaptation may occur during early stages of the disease, to compensate for ongoing neural dopamine death. Identifying these patterns and one day using them to facilitate early diagnosis first requires experimental protocols that can reliably capture subtle and transient locomotor changes over time.

Although traditional gait analysis in PD has historically relied on two approaches, using discrete markers or sensors to facilitate motion tracking, and manual review of video recordings ^[13,14]^, both methods present limitations. They can introduce observer bias and may lack the sensitivity required to consistently capture subtle or transient locomotor patterns, particularly those relevant in early disease stages. Recently, however, the field has increasingly adopted automated, markerless pose estimation methods, such as DeepLabCut (DLC). Early reports suggest that these can provide greater sensitivity to subtle gait changes ^[15–17]^. DLC provides automated tracking that minimizes observer bias and increases objectivity in data collection. At the same time, its fine temporal and spatial resolution enables the detection of subtle motor features that are often missed by conventional methods, making it well suited for preclinical PD studies.

This study aimed to characterize early gait alterations following partial dopaminergic lesions induced by low-dose 6-OHDA in the substantia nigra pars compacta (SNc) of Wistar rats. Locomotor behavior was tracked over six weeks to examine the influence of lesion extent and sex on gait parameters. To achieve this, we employed markerless pose estimation with DeepLabCut (DLC), which provides fine-grained kinematic resolution. This approach improves detection of subtle, sex-dependent motor alterations, enhancing the translational value of rodent models for prodromal Parkinson’s disease.

## MATERIALS AND METHODS

### Animals

Subjects were 102 Wistar rats (*Rattus norvegicus*), 58 males and 44 females, were bred at the Neuroscience and Behavior Laboratory at Universidad de Los Andes in Bogotá, Colombia. Subject ages were between postnatal days (PND) 60 and 80 at the start of the experiment. Subjects were randomly paired in standard laboratory cages measuring 16.5 × 50 × 35 cm. Water and food were provided ad libitum during the experiment. Environmental conditions were consistent throughout the study: a 12-12 hour light/dark cycle (lights on at 14:00), a temperature of 22 ± 2 °C, and a relative humidity of 57 ± 10%. Experiments were conducted during the dark portion of the cycle. The experimental protocol was approved by and followed the guidelines established by the Institutional Committee for the Care and Use of Laboratory Animals (CICUAL) at Universidad de Los Andes (Approval number CICUAL_20_012, issue date May 18, 2020), and adhered to the national ethical regulations for animal research in Colombia (Law 84 of 1989 and Resolution 8430 of 1993, Ministry of Health).

### Surgery and 6-OHDA lesion

Subjects were randomly assigned to receive either 6-hydroxydopamine (6-OHDA) or a vehicle solution. Rats were anesthetized with Isoflurane, craniotomies were created, and a 27-gauge needle was inserted unilaterally in the substantia nigra compacta (SNc) following the stereotaxic coordinates by Paxinos and Watson (2007): AP –5.25 mm, ML 2.2 mm, and DV 7.8 mm from Bregma. 6-OHDA solution (Sigma-Aldrich) was prepared at a concentration of 10 μg/μl in 0.9% saline contained ascorbic acid 0.02% w/v (≈ 0.2 mg/mL). A total volume of 0.25 μl (2.5 µg) was infused into the SNc via a 10 μl Hamilton™ syringe linked to a NE-1000 microinjection system (New Era Pump Systems Inc.) at a rate of 0.125 μl/min (infusion duration ≈ 2 min). After infusion, the needle was left in place for 5 additional minutes to allow diffusion. Control subjects received an equal volume of 0.9% Vehicle. Postoperative management involved 3-5 days of Meloxicam® administered subcutaneously.

### Experimental design

One week prior to surgery, all subjects were handled daily for 10 minutes to facilitate habituation. On day six, baseline motor performance was assessed using the horizontal ladder task. Surgeries were performed the following day, after which animals were randomly assigned to one of three groups, differing by the duration of their post-lesion motor evaluation interval: two weeks (t1), four weeks (t2), or six weeks (t3) (Fig 1A).

**Fig 1.**
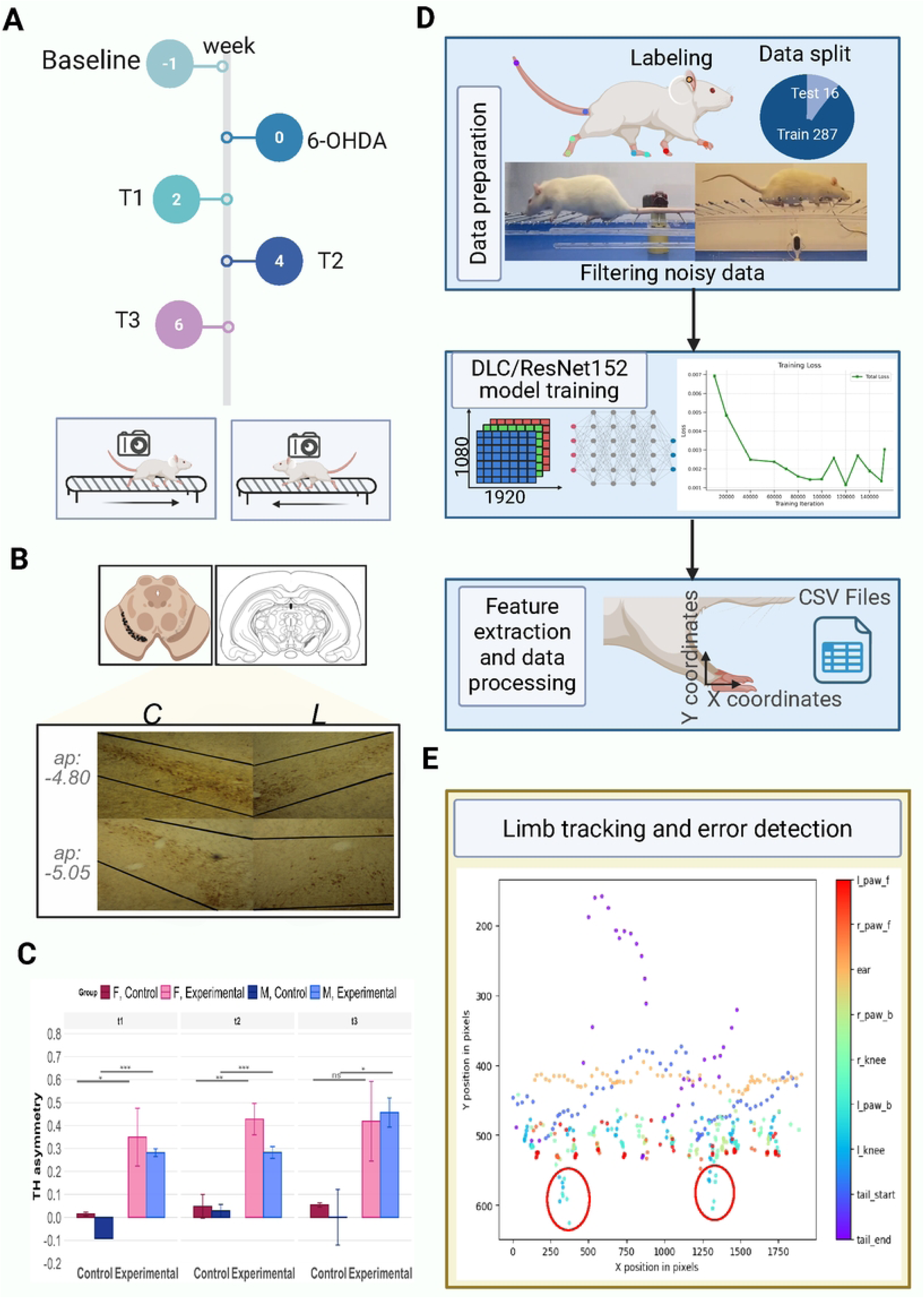
Schematic overview of the workflow applied to horizontal ladder walking recordings for the extraction of footfall errors and velocity measurements. **A.** *Data acquisition*: Videos were recorded using a Nikon camera positioned laterally to the ladder, capturing the right side of the rat during each trial. Recordings were collected at Baseline (−1), and up to two (t1), four (t2) or six (t3) weeks after surgery for each group, respectively. **B.** Relative TH-reactivity asymmetry showing the SNc coordinates. **C.** EMMs ± 95% CIs by Sex × Treatment × Time. Bars show EMMs (HC3-based 95% CIs) from the main-effects GLM. Brackets indicate E vs C contrasts within each Sex × Time with Holm-adjusted significance (*p* < .05 = *, < .01 = **, < .001 = ***, “ns” = not significant). Estimates and CIs correspond to those reported in Table 3. Sex: females (F) and males (M); Treatment: Control and Experimental (lesioned); Time: week 2 (t1), week 4 (t2), and week 6 (t3). **D.** *AI model design*: A total of 303 frames were manually labeled with 9 anatomical keypoints. These were split into 287 frames for training and 16 for testing. A pose estimation model was trained using DLC with the ResNet-152 architecture. The resulting X and Y coordinates for each body part were exported as .csv files. **E.** *Database creation and analysis*: Two behavioral metrics were extracted from the dataset: (1) the number of footfall errors, defined as displacements exceeding a paw-specific threshold and occurring in frames with a likelihood score > 0.75, and (2) the velocity in the X-direction, calculated using the displacement of the tail base across frames and reported in pixels per second (px/s).

### Gait measurements in the horizontal ladder

The structure of the horizontal ladder consisted of a transparent acrylic corridor measuring 100 × 15 × 62 cm (L × W × H), with steel rungs spaced 4 cm apart (Fig 1A). At one end of the ladder, a home cage was connected via a short ramp, allowing subjects to transition smoothly from the ladder into their familiar environment. At the beginning of each trial, rats were placed at the opposite end of the ladder and were expected to traverse the entire length without stopping, thereby initiating the test voluntarily. Trials ended when the animal reached the home cage, with no time limit imposed; if an animal paused, gentle tactile stimulation of the back was used to prompt movement. Each subject completed up to three trials per weekly session, separated by 2-minute intervals to minimize habituation. The trial with the fewest stops was selected for analysis, and when a subject completed the first crossing without interruption, that trial was used. Gait performance was quantified by measuring two variables: running velocity, calculated from the time the four paws first contacted the rungs until the head reached the end of the ladder; and footfall errors, defined as instances in which a paw slipped off a rung. Each trial was video-recorded individually and subsequently categorized for blinded analysis.

### Data Acquisition

Video recordings were obtained using either an iPhone X and a Nikon Coolpix B500 camera positioned laterally and orthogonal to the horizontal ladder at a fixed distance to maintain a consistent field of view. Early recordings captured with a fisheye lens were excluded from the training dataset to prevent distortion-related errors. High-definition videos (1920 × 1080 pixels) were recorded at 30 fps, providing a right-side lateral view of locomotor activity (Fi 1D). A total of 303 frames (287 for training, 16 for testing) were manually selected from sixteen videos to ensure representative coverage of postures and paw movements, including running, standing, missteps, and hopping. Nine anatomical landmarks were annotated using the DeepLabCut (DLC) graphical user interface (GUI): left and right hind paw joints (ankle, knee), both forepaws, tail base and tip, and right ear (Fig 2A). All frames preserved the original resolution and orientation; no cropping or downsampling was applied.

**Fig 2.**
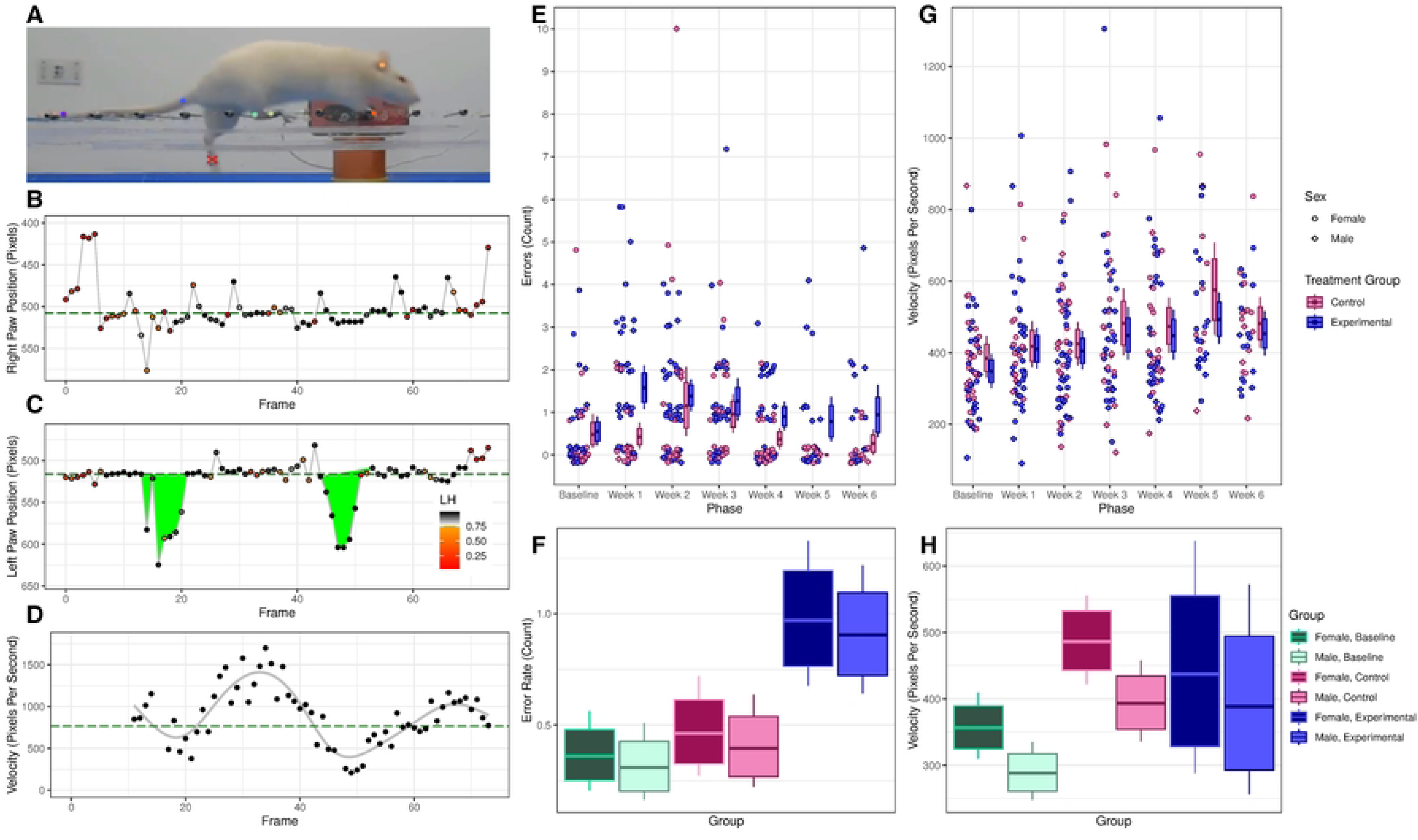
Analysis of step errors and velocity. **A.** Estimated anatomical tags overlaid on a representative video frame, according to DeepLabCut. The red X is a manually classified footfall error. **B.** Estimated horizontal position of the back right paw in each frame of a representative video. Points are color-coded according to each estimate’s likelihood. Only grayscale points (LH > .75) were “confirmed” and used for classification. No footfalls with this paw were classified as errors. **C**. As in panel B, for the back left paw. Two footfall errors were detected, filled in green. Errors began with the first confirmed position before threshold, and ended with the first confirmed position above baseline. **D.** Estimated interpolated velocity for confirmed foot positions. Also shown is a representative curve (generalized additive model, solid gray line), and overall mean velocity (total distance/total time, dashed green line). **E.** Footfall errors (circles = females, diamonds = males) per week by the control (pink) and experimental (blue) groups. Bootstrapped means are reported (bars = 80% confidence intervals, whiskers = 95% confidence intervals). Points represent discrete counts, but were randomly jittered on both axes to assist visual inspection. **F.** Summary of estimated error rates for an average subject in each condition (green = baseline, pink = control, blue = experimental), for both sexes (darker colors = female, lighter colors = male). Posterior estimates were computed using Equations 1 and 2, limited to only fixed effects (all *β* terms) and the population mean across subjects (*η*). Estimates for the control and experimental conditions here averaged performance across all six weeks (bars = 80% posterior credible intervals, whiskers = 95% credible intervals). **G.** As in panel E, but plotting overall velocities (pixels/second). Points were jittered on the horizontal axis to assist visual inspection. **H.** As in panel F, but plotting overall velocities (pixels/second).

### Immunohistochemistry

At the conclusion of each subject’s motor evaluation interval, they were euthanized by an overdose of sodium pentobarbital (85 mg/kg, i.p.) and transcardially perfused with 100–150 ml of 0.1 M phosphate-buffered saline (PBS) followed by 100–150 ml of 4% paraformaldehyde (PFA) for tissue fixation. Brains were extracted and post-fixed in PFA at 4°C. Tissue slicing (20 μm thick) was conducted using the Compresstome® VF-300-0Z microtome and then stored in Eppendorf tubes with 0.1 M PBS at 4°C for further processing.

Immunohistochemistry was carried out according to the guidelines provided by Abcam for the Anti-Tyrosine Hydroxylase antibody (Abcam ab6211) and the Rabbit Specific HRP/DAB Detection IHC Kit (Abcam ab64261). Tissue samples were placed on glass slides and rinsed twice with 0.1 M PBS for a duration of 10 minutes. To inhibit endogenous peroxidase activity, a hydrogen peroxide blocking solution was used for 10 minutes at 15–18°C, followed by three washes with PBS. Subsequently, a protein blocking solution was applied for 10 minutes and removed with two more washes of PBS. The primary antibody (Anti-Tyrosine Hydroxylase, diluted 1:1000) was introduced and incubated overnight at 4°C. The next day, sections were rinsed four times with PBS and incubated with the secondary antibody solution for 20–30 minutes. After four more washes, the chromogenic substrate diaminobenzidine (DAB) was added for 1–10 minutes to detect immunoreactivity, followed by three final washes with PBS. Tissues were finally mounted on glass slides and covered with coverslips secured using Canada balsam. Images were captured using a Canon EOS Rebel T3i camera that was attached to a Meiji MT5000 microscope (10× objective). Photomicrographs were obtained from both sides of the SN separately.

#### Statistical analyses: Video Analyses

Pose estimation was performed using DLC versions 2.3.10 and 2.3.11 with a ResNet-152 backbone pre-trained on ImageNet (Fig 1D). This architecture was selected over lighter alternatives due to the presence of motion blur, rapid limb movements, and the limited dataset size, which favored a more expressive feature extractor. The network was trained for 160,000 iterations using stochastic gradient descent. The effective batch size was set at 4 due to GPU memory constraints. Training was conducted on Google Colab with NVIDIA T4 and A100 GPUs (16 GB and 40 GB VRAM, respectively), running Python 3.11 and CUDA 11.8. Model accuracy was quantified as mean average error (MAE) in pixels for both training and test sets. After training and validation, the DLC model was applied to the remaining video dataset. For each frame, X–Y coordinates were extracted for all annotated landmarks (Fig 1D). Frames with a likelihood score below 0.75 were excluded from further analysis.

Footfall errors were independently calculated for each paw by determining a vertical baseline from a trimmed mean of Y-coordinates. A candidate error was defined as beginning with a downward excursion beyond an adaptive threshold, calculated as the paw’s baseline plus a fixed offset (35 pixels in our implementation). This adaptive approach ensured that thresholds were referenced to each paw’s height, accounting for differences in limb perspectives across recordings ^[16,18]^. This error footfall was then considered to continue until the first foot position confirmed (LH > 0.75) to be above baseline. Consolidating all such frames into a single error epoch prevented double-counting of transient fluctuations during the same footfall (Fig 2B and C). For each footfall error, maximum drop distance and epoch duration (frames) were recorded.

Horizontal velocity was estimated using the X-coordinate of the tail base landmark, which showed the lowest incidence of occlusion (Fig 2D). Average velocity (pixels/s) was calculated for the first half, second half, and total duration of each trial. Time was reconstructed from frame counts assuming a constant 30 fps. Due to the absence of spatial calibration, velocities are expressed in pixels rather than converted to physical units.

### Behavioral analyses

Three behavioral measures were assessed (total footfall errors, proportion of left-side footfall errors, and velocity) using multi-level Bayesian regression models, implemented using the Stan programming language ^[19]^. All models used the same overall structure, characterizing performance for a given subject s during a particular week was a sum of fixed and random effects, given by *μ*_s,w_:

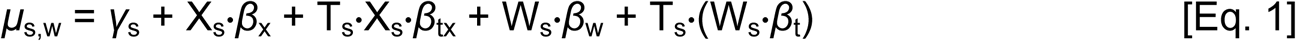

Here, *γ_s_* denotes a random intercept specific to subject *s* that was constant across the experiment. X*_s_*denotes a centered dummy variable for the subject’s sex (female = -0.5, male = 0.5) and T*_s_* denotes a dummy variable for treatment (control = 0, experimental = 1). *β_x_* thus denotes a fixed effect of sex and *β_tx_* denotes a sex-by treatment interaction term. **W***_s_* is a vector of six dummy values specifying which week was being modeled during the experiment; only one value in **W***_s_*was set to 1 at a time, with the rest set to zero. To measure baseline performance, all six values in **W***_s_* were set to zero. Finally, **β***_w_* denotes a vector of six fixed effects for each week of the experiment (constituting that week’s difference from baseline performance), and **β***_t_* denotes six fixed effects of treatment during each week (constituting the difference between experimental and control conditions during that week). Thus, in total, performance was modeled using one random parameter per subject, as well as 14 fixed effects.

Using the structure described above, total errors were modeled as a Poisson process governed by a rate *θ*. Model parameters were estimated using regularizing priors:

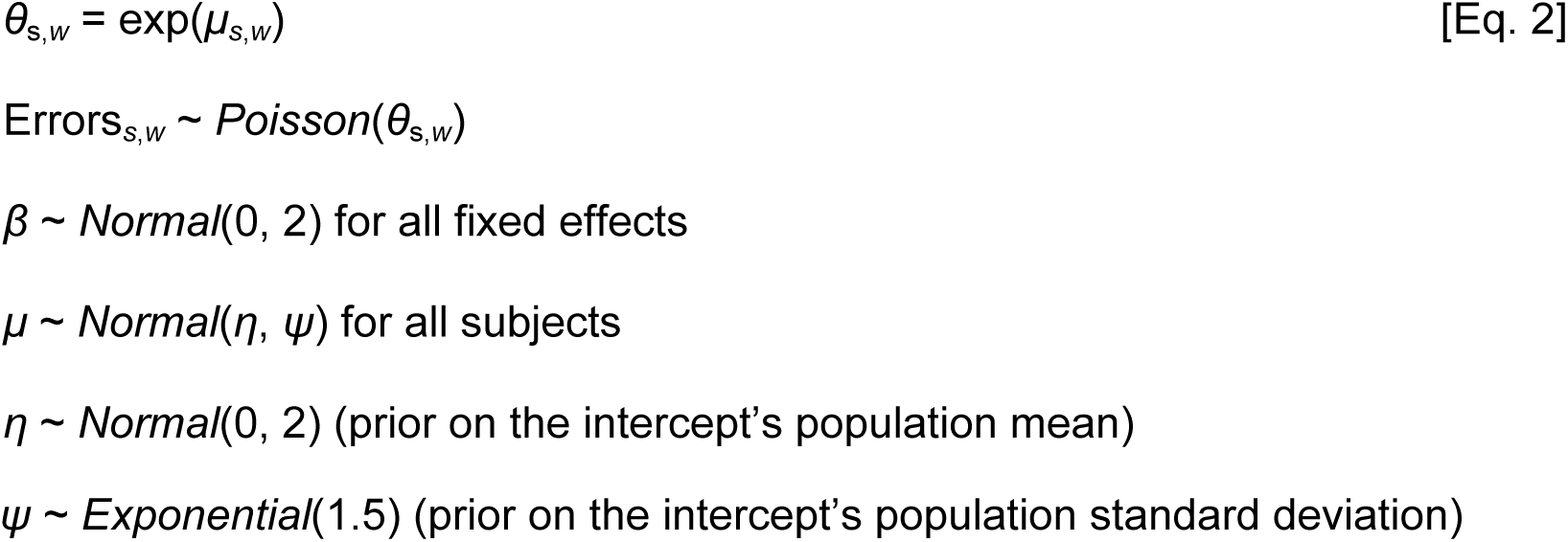

The asymmetry of errors was modeled by the total errors made on the subject’s left side as a binomial process, governed by a probability of left error *τ* for each error made. These were also estimated using the same structure to *μ_s_*_,*w*_, as described above (albeit different actual values for those parameters). The same regularizing priors were used as those in the Errors model.

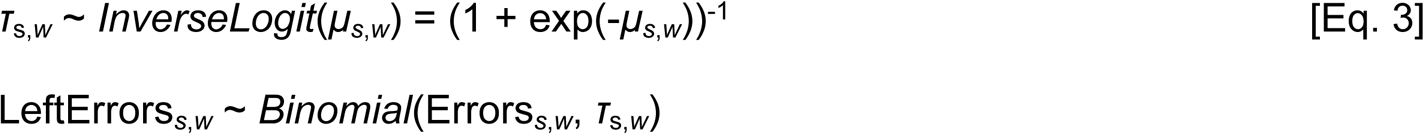

Finally, the velocity was modeled using a linear model after converting velocities to log units. This helped to manage the considerable positive skew of the velocity data. Residual error on log units was also treated as a fixed effect across subjects.

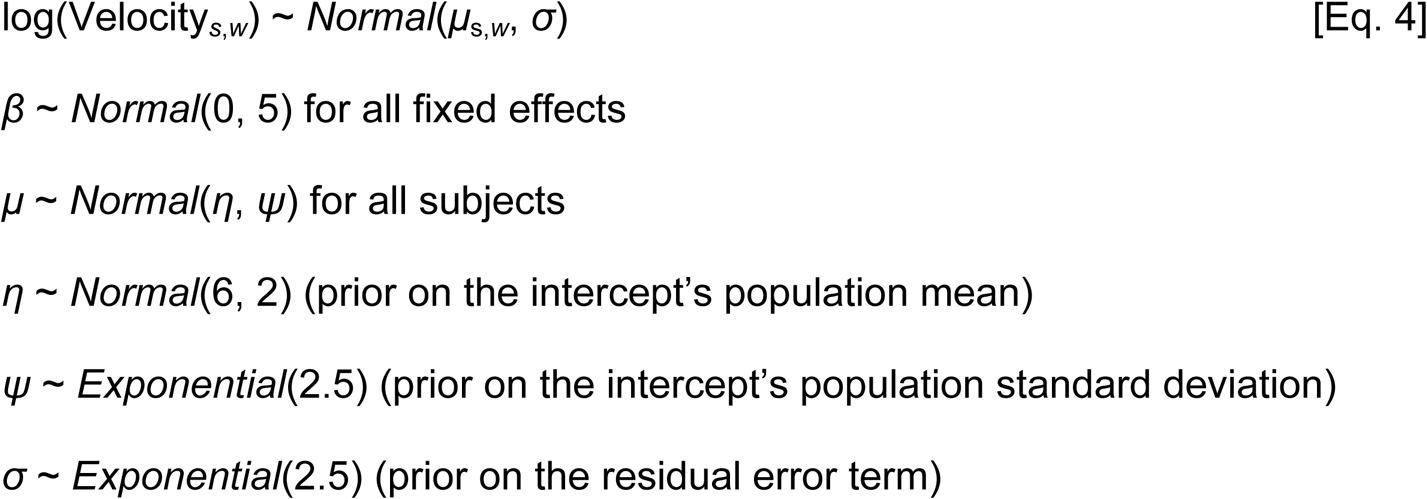

For each of the three models above, parameters were fit using Hamiltonian Monte Carlo, allowing numerical estimation of the uncertainty of both individual parameters and of derived values (such as a subject’s expected performance during any given week). All subjects were included in a single model, contributing data from those weeks in which their performance was recorded.

### Tyrosine Hydroxylase

A Gaussian generalized linear model with identity link was used to analyze asymmetry in tyrosine hydroxylase (TH)-positive neurons between hemispheres of the SNc. Statistical analyses were conducted in R (version 3.6.0), with categorical predictors coded as factors and Female (F), Control (C), and week (t1) set as reference levels. Analyses were restricted to complete cases (N = 43). To account for potential heteroscedasticity and unbalanced group sizes, heteroskedasticity-consistent (HC3) standard errors were applied, with significance set at α = .05. Sensitivity analyses included bias-corrected bootstrap confidence intervals (1,000 resamples) and cluster-robust standard errors by subject. Estimated marginal means (EMMs) were obtained for each Sex × Treatment × Time cell, and planned contrasts (Experimental vs. Control) were adjusted using Holm’s procedure.

### Data and Code Accessibility

All data supporting the findings of this study are included in the manuscript and are available from the corresponding author upon request.

## RESULTS

### DLC model performance and application to behavioral dataset

The ResNet-152 network trained on 303 curated frames achieved a test set MAE of 8.41 px without thresholding, which improved to 5.42 px with a p-cutoff ≥ 0.75. Training MAE was 3.80 px without the cutoff and 3.32 px with it. These results confirmed high pose estimation accuracy suitable for downstream gait analysis (Fig 2E). The trained model was then applied to the complete set of experimental videos. Error analysis revealed limb-specific deviations consistent with missteps, meeting all three detection criteria (baseline deviation, high confidence, and single-event consolidation). Average velocity was calculated for the first half, second half, and entire trial, allowing temporal segmentation of locomotor performance. Taken together, these outputs provided the basis for statistical analyses evaluating differences between lesioned and control groups throughout the study.

### Behavioral Results

There was considerable variation both between subjects and across sessions, as illustrated in Figure 2E, which plots the errors made by each subject during each session. The modal outcome for any given phase of the experiment was zero errors, with considerable positive skew. Bootstrapped means per condition suggest that 0.5 to 1.0 errors per session are a typical rate. The corresponding traversal velocities also showed some positive skew and suggested a gradual acceleration across weeks, as illustrated in Figure 2G.

In order to estimate the effects in a manner that controls for covariates, parameters were fit for the models described in Equations 1-4. Overall performance for each of the conditions (pre-treatment, control, and experimental), averaged across all weeks, given an average subject of each sex, is summarized in Figure 2F. At baseline, an average subject was expected to make 0.331 errors [CI_95%_: 0.208, 0.491], compared to 0.425 errors [CI_95%_: 0.277, 0.605] in the control condition and 0.934 errors [CI_95%_: 0.715, 1.184] in the experimental condition. These results indicate that the experimental treatment resulted in about 0.508 additional errors [CI_95%_: 0.255, 0.776] than did the control treatment. However, a reliable difference in mean errors did not arise as a function of sex. Figure 2H depicts a similar summary for velocity. This shows that shows that, at baseline, an average subject was expected to traverse the apparatus at 320.7 pixels per second [CI_95%_: 290.5, 352.6], compared to 436.2 pixels per second [CI_95%_: 392.5, 481.7] in the control condition and 405.4 pixels per second [CI_95%_: 370.7, 442.5] in the experimental condition. Thus, overall, subjects in both conditions completed the traversal faster during the experimental phases than they did during the initial baseline measurement.

### Footfall errors

Posterior predictions for error rates for an average rat of each sex in all phases of the experiment are plotted in Figure 3A. These are computed based on the model’s fixed effects, with an intercept equal to the population average. These parameters, along with their 95% posterior credible intervals are shown in Table 1. Overall, errors in the experimental group rise immediately following the experimental treatment, and remain elevated for the duration of the experiment. Error rates also rise in the control group, but only during weeks 2 and 3 of the study, falling back to or even below baseline levels in weeks 4 through 6. No reliable difference is observed due to sex.

**Fig 3.**
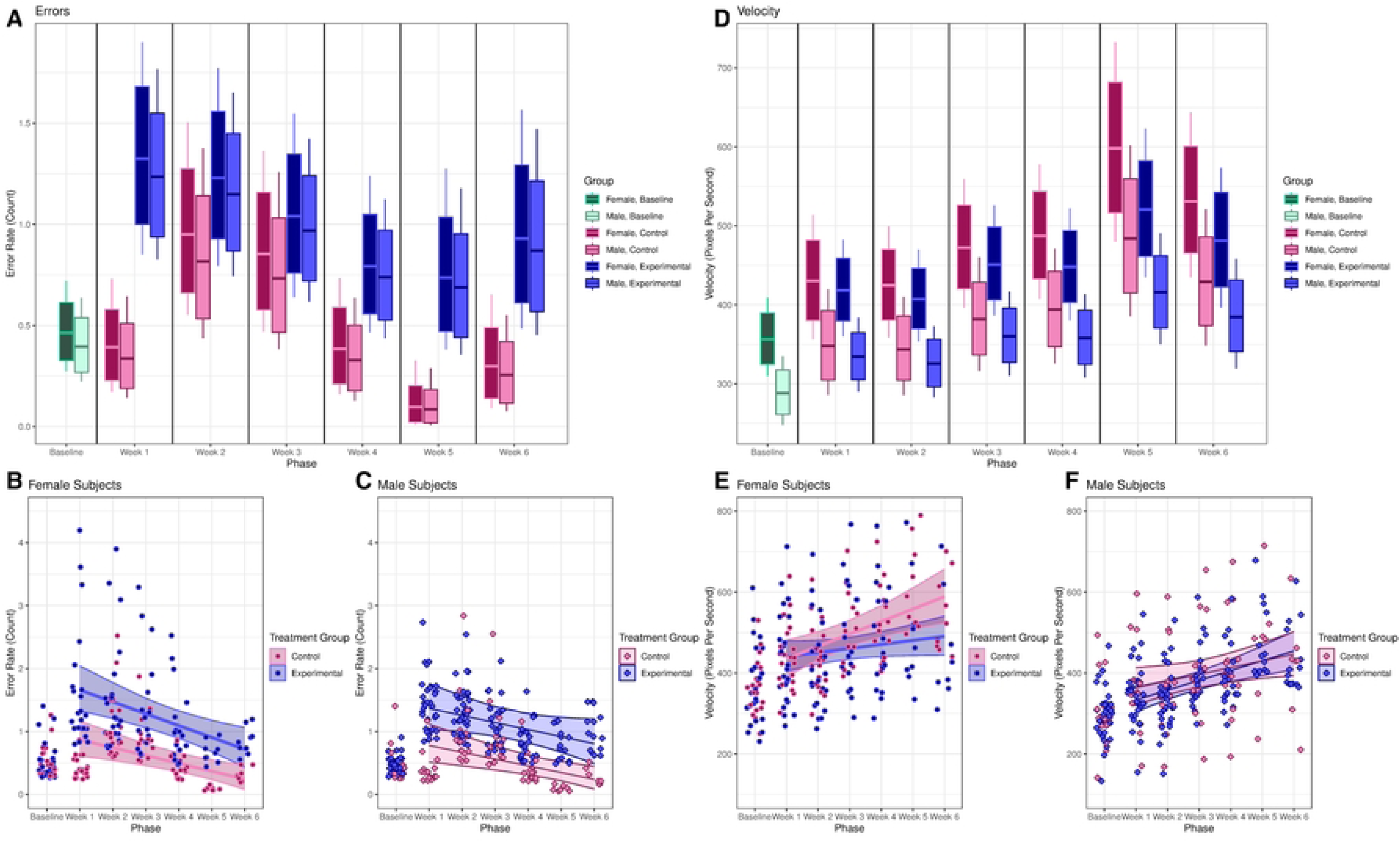
Week-by-week estimates of performance. **A.** Estimated error rate for an average subject during each week, split by condition (green = baseline, pink = control, blue = experimental) and sex (darker colors = female, lighter colors = male), based on the population-level effects in Equations 1 and 2 (bars = 80% posterior credible interval, whiskers = 95%). **B.** Posterior mean estimates of individual error rates per session for each female subject, based on the full model in Equations 1 and 2. Each point reflects one subject during that session. Subjects are only plotted for weeks where their data are available. Trendlines show the overall linear trend across subjects for the weeks after baseline (shaded region = 95% credible interval). **C.** As in panel C, but plotting male subjects. D-F. As in panels A-C, but plotting estimated velocities (pixels/second).

**Table 1.** Bayesian Model Parameters for Errors (log units). * Bolded/shaded rows are fixed effects whose posterior credible intervals exclude zero. Green rows consistently make positive contributions, whereas red rows consistently make negative ones.

| Effect | Scale | Parameter | Mean | 95% CI |
| --- | --- | --- | --- | --- |
| Pop. Intercept Mean | log(error rate) | $\eta$ | -0.875 | [-1.284, -0.502] |
| Pop. Intercept SD | log(error rate) | $\psi$ | 0.611 | [0.433, 0.817] |
| Sex | Female = -0.5,<br>Male = 0.5 | $\beta_x$ | -0.160 | [-0.783, 0.464] |
| Sex × Experiment | Interaction | $\beta_{tx}$ | 0.092 | [-0.662, 0.878] |
| Week 1 | Dummy Code | $\beta_w[1]$ | -0.210 | [-1.007, 0.496] |
| **Week 2** | ~ | $\beta_w[2]$ | **\*0.707** | **[0.150, 1.262]** |
| **Week 3** | ~ | $\beta_w[3]$ | **\*0.597** | **[0.005, 1.188]** |
| Week 4 | ~ | $\beta_w[4]$ | -0.240 | [-1.068, 0.526] |
| **Week 5** | ~ | $\beta_w[5]$ | **\*-1.849** | **[-3.716, -0.372]** |
| Week 6 | ~ | $\beta_w[6]$ | -0.527 | [-1.609, 0.455] |
| **Week 1×Experiment** | **Interaction** | $\beta_t[1]$ | **\*1.313** | **[0.597, 2.099]** |
| Week 2×Experiment | ~ | $\beta_t[2]$ | 0.322 | [-0.220, 0.853] |
| Week 3×Experiment | ~ | $\beta_t[3]$ | 0.263 | [-0.342, 0.887] |
| **Week 4×Experiment** | ~ | $\beta_t[4]$ | **\*0.820** | **[0.014, 1.698]** |
| **Week 5×Experiment** | ~ | $\beta_t[5]$ | **\*2.340** | **[0.826, 4.202]** |
| **Week 6×Experiment** | ~ | $\beta_t[6]$ | **\*1.254** | **[0.191, 2.381]** |

The estimated error rates for each subject during every phase of the study in which they appeared are displayed in Figure 3B (females) and 3C (males). As such, estimates for all subjects appear at baseline and in weeks 1 and 2, but later weeks only include estimates for those cohorts still contributing data during those phases. Although the statistical model treated the six post-treatment phases as nominal variables for maximum flexibility, we include a post-hoc estimate of the mean linear trend throughout the post-treatment phases for each group and each sex. In general, errors tended to fall over the course of the experiment: Post-hoc linear trends of change in errors per week consistently had negative slopes in most groups (control females: -0.123 [CI_95%_: -0.202, -0.053]; experimental females: -0.188 [CI_95%_: -0.292, -0.084]; control males: -0.106 [CI_95%_: -0.181, -0.038]), with the exception of experimental males (−0.112 [CI_95%_: -0.223, 0.003]). Overall, this reveals that although females overall had similar means to males, experimental females appeared slightly more variable, with larger outliers during the first few post-treatment weeks. Nevertheless, these results indicate that the variation in error rates was chiefly driven by a difference between the control and experimental conditions, without a consistent effect attributable to sex.

### Asymmetry analyses

An additional analysis evaluated whether error frequency differed between left and right paws (Equation 3). Subjects overall made consistently more errors with their left paws (74.0% [CI_95%_: 57.5%, 87.1%]). However, due to the low error rates overall, no other consistent differences could be detected, whether as a function of sex, treatment or phase. Independent cross-check verification of the original video recordings and the corresponding DLC-extracted data confirmed that the observed asymmetry was not attributable to differences in the software’s detection of left-versus right-paw errors, nor to misclassifications.

### Velocity Analyses

Posterior predictions of average velocities for males and females in all experimental phases are presented in Figure 3D, based on the fixed effects and a population-mean intercept. Model parameters, along with their 95% posterior credible intervals, are provided in Table 2. Overall, subjects consistently complete the task more quickly following treatment than they did at baseline, and females traverse the apparatus more rapidly than males (including at baseline). No consistent treatment effect was observed, since weekly control group velocities closely matched those of the experimental group.

**Table 2.**
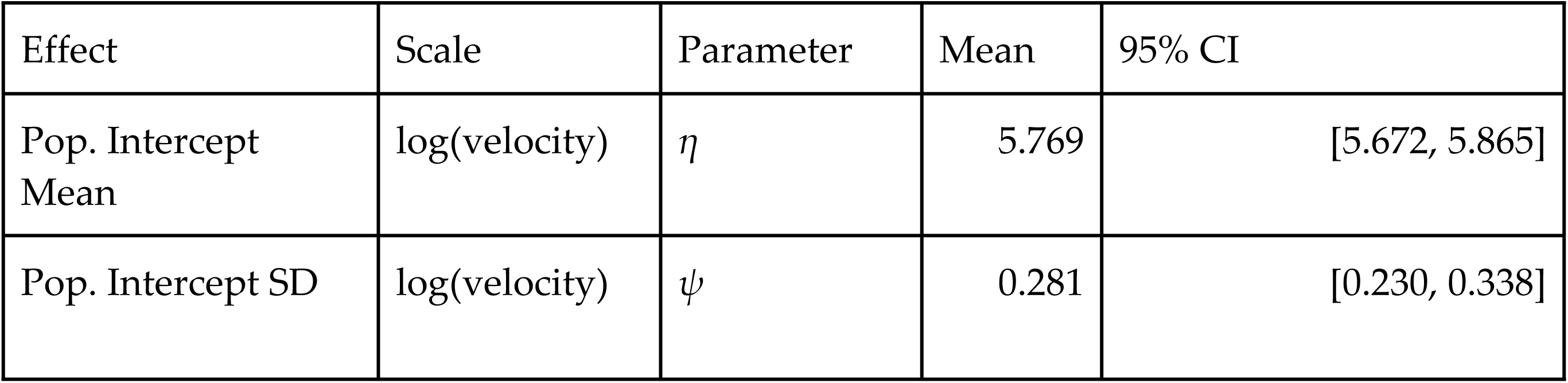
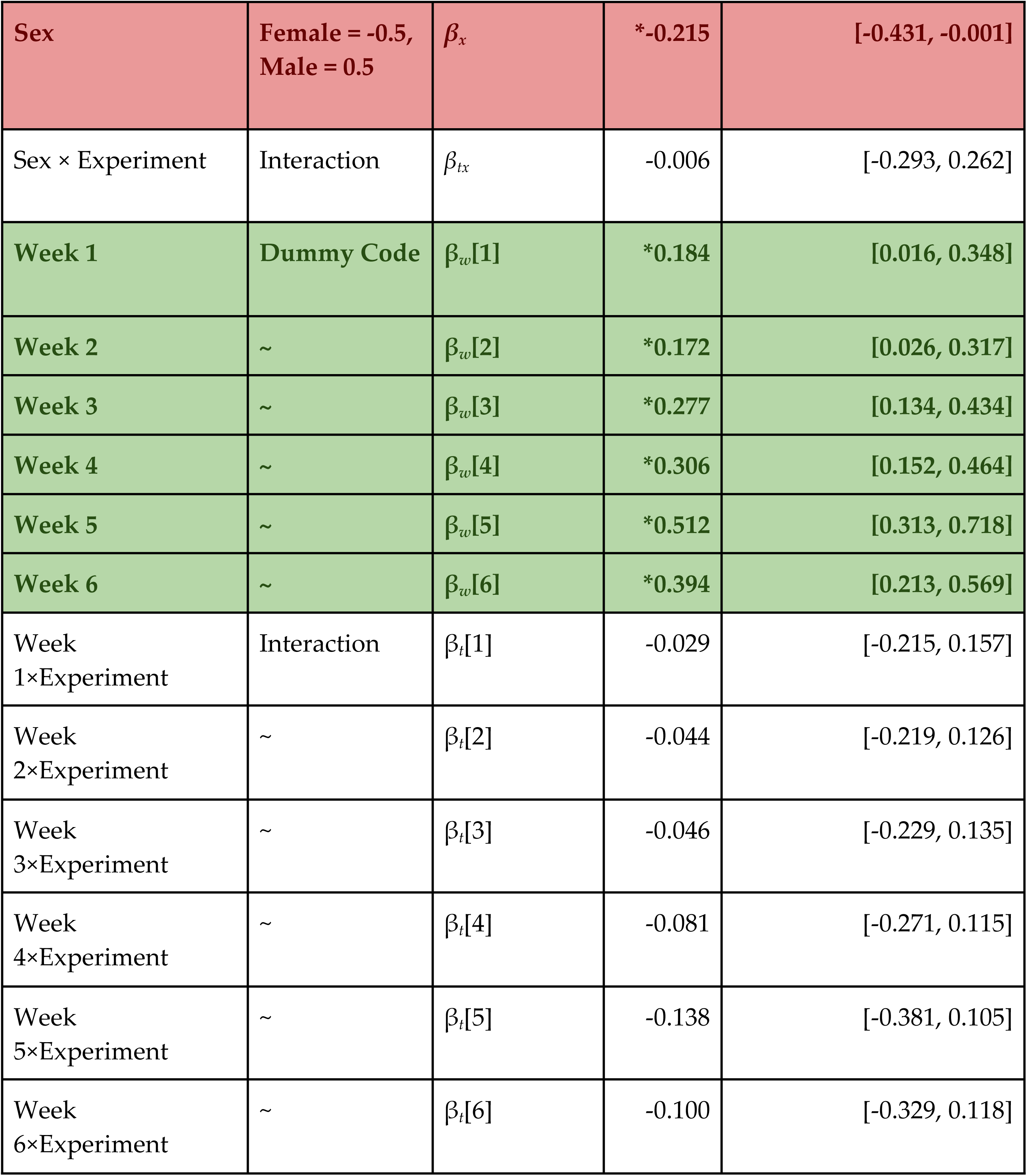
Bayesian Model Parameters for Velocity (log units). * Bolded/shaded rows are fixed effects whose posterior credible intervals exclude zero. Green rows consistently make positive contributions, whereas red rows consistently make negative ones.

Estimated velocities for each subject across all phases of the study in which they appeared are shown in Figure 3E (female subjects) and 3F (male subjects), using the same post-hoc linear analysis described for Figure 3B & 3C. Once the post-treatment weeks are treated as ordered variables, a striking sex-by-treatment interaction emerges. Control females begin fast and also accelerate, their velocity increasing by 30.6 [CI_95%_: 12.7, 48.8] pixels/second each week, whereas experimental females did not consistently accelerate (a 10.1 [CI_95%_: -2.4, 23.6] pixels/second change each week). Males, on the other hand, were generally more consistent: Both control males (16.5 [CI_95%_: 1.36, 32.4] pixels/second each week) and experimental males (24.2 [CI_95%_: 12.4, 36.7] pixels/second each week) increased their velocity over time. This pattern suggests that while male trajectories were uniform across groups, female performance was more sensitive to lesion status, highlighting potential sex-specific strategies of motor adaptation.

### Tyrosine Hydroxylase

The analysis revealed a robust main effect of treatment, with Experimental animals showing greater TH-reactivity asymmetry than Controls (Table 3). Effects of time were modest, with only t3 showing a small increase relative to t1, and the main effect of sex was not reliable once treatment and time were considered. Planned contrasts further confirmed higher asymmetry in the Experimental group across most Sex × Time cells. At two weeks (t1), differences were evident in both females (Δ = 0.33, p = .038) and males (Δ = 0.37, p < .001). At week 4 (t2), Experimental animals again exceeded Controls in females (Δ = 0.38, p = .0014) and males (Δ = 0.25, p < .001). By week 6 (t3), the pattern persisted in males (Δ = 0.46, p = .019), while the female comparison (Δ = 0.36, p = .097) pointed in the same direction but remained inconclusive due to wider confidence intervals. Despite this reduced precision, the overall pattern Experimental > Control was consistent across sexes and time points (Figure 1, panels B and C). Importantly, results were robust to alternative specifications, as conclusions were unchanged when bootstrap intervals, cluster-robust standard errors, or exploratory mixed-effects models were applied.

**Table 3.** Estimated marginal means (EMMs) of relative TH-reactivity asymmetry by Sex × Treatment × Time, and Experimental–Control (E−C) contrasts. Values are EMMs ± 95% CIs from a Gaussian GLM with identity link and main effects of sex, treatment, and time; inference uses heteroskedasticity-consistent (HC3) standard errors. Planned contrasts compare E vs C within each Sex × Time cell with Holm adjustment for six tests (two-sided α = .05). Positive differences indicate higher asymmetry in the Experimental group.

| Sex | Time | Control mean ± 95% CI | Experimental mean ± 95% CI | Diff. (E – C) | p (adjusted) |
| --- | --- | --- | --- | --- | --- |
| F | t1 | 0.015 ± [-0.004, 0.034] | 0.350 ± [0.036, 0.663] | 0.334 | 0.0378 |
| M | t1 | -0.091 ± [-0.091, -0.091] | 0.282 ± [0.240, 0.324] | 0.373 | < .0001 |
| F | t2 | 0.048 ± [-0.102, 0.198] | 0.428 ± [0.266, 0.589] | 0.38 | 0.0014 |
| M | t2 | 0.028 ± [-0.042, 0.098] | 0.282 ± [0.224, 0.341] | 0.254 | < .0001 |
| F | t3 | 0.054 ± [0.034, 0.075] | 0.419 ± [-0.014, 0.852] | 0.364 | 0.0965 |
| M | t3 | 0.001 ± [-0.348, 0.350] | 0.456 ± [0.317, 0.596] | 0.456 | 0.0191 |

## DISCUSSION

The present study examined gait alterations in Wistar rats with partial unilateral dopaminergic lesions induced by 6-OHDA in the SNc, a model designed to approximate prodromal Parkinson’s disease. Running velocity and footfall errors were tracked over six weeks to capture dynamic motor changes that reflect early dopaminergic disruption. Lesioned animals showed greater TH-reactivity asymmetry than controls, and this histological difference was reflected in persistent gait inaccuracies. In terms of locomotor velocity, both lesioned and control males displayed similar, gradual increases across weeks, with no significant differences between groups. By contrast, control females showed a marked acceleration, whereas lesioned females exhibited less pronounced increases, resulting in greater variability. This divergence suggests that partial dopaminergic depletion in the SNc produces enduring, sex-specific motor alterations, expressed as gait deficits that remain detectable but evolve in severity over time, and distinct velocity trajectories across weeks and sexes.

In contrast to the pronounced and progressive motor decline typically observed in high-dose 6-OHDA models ^[11,20]^, the partial dopaminergic lesions in our study produced a more nuanced profile of gait alteration. Across the six weeks of testing, deficits were subtle yet sustained, showed partial reversibility, and were expressed primarily in precision and sex-specific variability in velocity. These findings align with clinical observations in early PD, where gait alterations may precede overt bradykinesia ^[21]^. Footfall errors in lesioned animals rose immediately after lesion, peaking at week three and remaining elevated thereafter. Although average error rates were similar between sexes, females in the lesioned group showed greater variability (Figure 3B) than males (Figure 3C), suggesting potential sex-specific differences in adaptation. Some of the variability observed could reflect sex-related differences, which might influence sensitivity to the 6-OHDA procedure. Importantly, analysis of TH reactivity confirmed consistent asymmetry across experimental groups and sexes, indicating that lesion extent was comparable. Thus, while subtle individual variability cannot be excluded, the observed effects are better interpreted as sex-differences in sensitivity to comparable dopaminergic depletion rather than uneven lesion induction. Furthermore, error rates in the control group increased transiently during weeks two and three before returning to baseline, whereas lesioned animals maintained higher error levels across sessions, underscoring lesion-specific effects beyond procedural influences ^[22–24]^. Taken together, our findings enabled the differentiation between transient procedural effects and sustained impairments attributable to partial SNc lesions, while also capturing sex-specific trajectories of motor adaptation.

Moreover, our findings add evidence that supports the viability of low-dose 6-OHDA lesions in the SNc as a model of prodromal motor impairment. Unlike high-dose protocols or mild lesions in the MFB or DS ^[25–27]^, which often produce more severe deficits, this approach reproduces subtle, stable alterations that might mirror early disease stages. By focusing on partial dopaminergic lesions, this model holds potential for identifying early behavioral markers and testing interventions aimed at slowing disease progression.

The observation that partial SNc-unilateral lesion resulted in subtle, dynamic, and partially reversible impairments aligns with the view that early-stage PD is not defined by uniform deterioration, but by a complex interplay between deficit and compensation ^[28,29]^. The slight reduction in velocity observed only in lesioned females, despite consistent increases in footfall errors in both sexes, suggests that partial lesions may differentially affect fine motor control rather than gross locomotor drive. In humans, similar dissociations between speed and gait accuracy have been reported in cohorts of early stages of PD identified via REM sleep behavior disorder or genetic risk ^[30–32]^, where stride variability increases without slowing gait. Such parallels strengthen the translational value of refining low-dose lesion protocols to better represent early motor phenotypes.

Sex differences refine this interpretation as females in the lesioned group showed greater variability in footfall errors and velocities, suggesting that female subjects engaged in a wider range of adaptation strategies. This pattern resonates with clinical evidence that women with PD often present later onset ^[33]^ but may display different compensatory recruitment patterns in cortico-striatal circuits^[34]^. Rodent models ^[35]^ and clinical studies ^[36]^ have suggested that estrogen may confer some neuroprotective effects and enhance dopaminergic plasticity. In our study, however, lesioned females displayed greater variability in velocity and accuracy, which could indicate that early-stage gait alterations in females follow a wider range of trajectories, possibly involving both earlier increased susceptibility to subtle impairments and a compensatory capacity that may help delay the manifestation of overt motor symptoms. These results underscore the importance of systematically including both sexes in preclinical PD research to elucidate the mechanisms underlying sex-specific trajectories of motor alteration and recovery in early Parkinson’s.

Although control animals also showed a post-surgical increase in errors, this rise was smaller, shorter in duration, and showed clear improvement from week three onward. In contrast, lesioned animals maintained elevated error rates across the study, despite partial recovery in later weeks. While we cannot exclude other contributing factors, such as habituation to the ladder, improved task efficiency, or stress and anxiety induced by exposure to a bright open setting, the proximity of the first assessments to the perioperative period may also have influenced performance. For example, anesthesia, postoperative inflammation, or analgesic treatment could have transiently affected gait, as suggested by the brief increase in errors among vehicle-treated controls. However, because all animals underwent the same procedures, the sustained and more frequent errors observed in lesioned subjects point to effects of dopaminergic depletion beyond those attributable to perioperative or learning-related influences. This evidence suggests that in early stages of dopamine depletion the distinctive features of gait dynamics may not depend solely on the magnitude of errors but rather on their timing, persistence, and patterns of resolution. These temporal profiles provide informative markers for refining the characterization of prodromal dopaminergic depletion and for identifying potential intervention windows in which preventive or compensatory strategies may be most effective, an essential step toward translating preclinical findings into clinical applications ^[37,38]^.

The use of markerless pose estimation via DeepLabCut (DLC) enabled high-resolution detection of gait changes while reducing observer bias and requiring only limited manual labeling. Importantly, accurate estimates of velocity and footfall events were obtained using consumer-grade video equipment, demonstrating that detailed phenotyping of rodent locomotion can be achieved cost-effectively without specialized motion capture systems ^[16,17,39]^. These advantages make DLC an enabling technology for refining early-stage PD models, particularly in settings where specialized motion capture systems are unavailable. By improving measurement sensitivity, such tools can strengthen the translational value of early-stage PD models.

Despite these promising results, several limitations should be considered. The present approach prioritized longitudinal tracking of velocity and footfall accuracy, which provided a precise temporal map of early motor changes but limited the detection of more complex interlimb coordination patterns. The use of only two hind paw markers restricted error classification to a single category, leaving finer distinctions in paw dynamics undetected. Expanding the set of anatomical landmarks could capture additional kinematic variables, such as step-to-step variability or digit articulation^[40,41]^, thereby broadening the scope of the model for both gross and fine motor control.

Data availability also imposed constraints. The video corpus was relatively small, and some recordings were excluded from behavioral analysis because they were used to train the DLC model. Combined with the progressive decline in sample size across weeks, this reduced coverage at later time points and limited statistical power for detecting delayed effects. Technical conditions further restricted resolution, as recordings relied on a single lateral angle and low frame rate, introducing motion blur and lowering tracking confidence during high-speed forelimb movements. Although these issues did not compromise detection of velocity or footfall errors, higher frame rates, multi-angle acquisition, and larger, independent datasets would enhance kinematic resolution. Collecting more minutes of recording per animal each week would also allow DLC-based automation to support richer quantification of subtle gait features and refine early motor phenotyping. Taken together, these considerations do not undermine the central findings but highlight feasible strategies—richer video corpora, expanded kinematic features, and increased sampling—that can strengthen the utility of partial-lesion models for studying prodromal PD and improve their translational value.

In summary, this study demonstrates that low-dose 6-OHDA lesions in the SNc can induce temporal, sex-dependent gait inaccuracies and locomotor velocity, capturing key features that might resemble early-stage PD. The study revealed temporal patterns of sex-specific adaptation in control and lesioned subjects, and the use of DLC enhanced the detection of fine-grained gait alterations. Together, these findings support the refinement of partial lesion of the SNc by using the 6-OHA model for studying prodromal PD and reinforce the importance of integrating sensitive behavioral tracking tools to improve their translational potential.

## Acknowledgements

We thank Yaneth Camargo and Yaneth Gómez for their support in animal care and for maintaining laboratory operations during the pandemic closure. We also thank the undergraduate and graduate students of the Neuroscience and Behavior Laboratory for their assistance with manual video editing and constructive feedback throughout the project.

## Data Availability Statement

All relevant data are included within the manuscript and its Supporting Information files. Upon acceptance, the full repository (including videos, databases, and the DLC model code) will be made publicly available in the Zenodo repository at 10.5281/zenodo.17518250

## Author Contributions

**Conceptualization:** Diego Lievano Parra, Fernando P Cardenas.

**Data curation:** Diego Lievano Parra, Juan D Garavito, Greg Jensen.

**Investigation:** Diego Lievano Parra.

**Methodology:** Diego Lievano Parra, Fernando P Cardenas.

**Project administration:** Fernando P Cardenas.

**Supervision:** Fernando P Cardenas.

**Writing – original draft:** Diego Lievano Parra, Juan D Garavito, Greg Jensen, Valeria V González, Fernando P Cardenas.

## Supporting information

**S1 Fig 1. Schematic overview of the workflow applied to horizontal ladder walking recordings for the extraction of footfall errors and velocity measurements. A.** *Data acquisition*: Videos were recorded using a Nikon camera positioned laterally to the ladder, capturing the right side of the rat during each trial. Recordings were collected at Baseline (−1), and up to two (t1), four (t2) or six (t3) weeks after surgery for each group, respectively. **B.** Relative TH-reactivity asymmetry showing the SNc coordinates. **C.** EMMs ± 95% CIs by Sex × Treatment × Time. Bars show EMMs (HC3-based 95% CIs) from the main-effects GLM. Brackets indicate E vs C contrasts within each Sex × Time with Holm-adjusted significance (*p* < .05 = *, < .01 = **, < .001 = ***, “ns” = not significant). Estimates and CIs correspond to those reported in Table 3. Sex: females (F) and males (M); Treatment: Control and Experimental (lesioned); Time: week 2 (t1), week 4 (t2), and week 6 (t3). **D.** *AI model design*: A total of 303 frames were manually labeled with 9 anatomical keypoints. These were split into 287 frames for training and 16 for testing. A pose estimation model was trained using DLC with the ResNet-152 architecture. The resulting X and Y coordinates for each body part were exported as .csv files. **E.** *Database creation and analysis*: Two behavioral metrics were extracted from the dataset: (1) the number of footfall errors, defined as displacements exceeding a paw-specific threshold and occurring in frames with a likelihood score > 0.75, and (2) the velocity in the X-direction, calculated using the displacement of the tail base across frames and reported in pixels per second (px/s).

**S2 Fig 2. Analysis of step errors and velocity**. **A.** Estimated anatomical tags overlaid on a representative video frame, according to DeepLabCut. The red X is a manually classified footfall error. **B.** Estimated horizontal position of the back right paw in each frame of a representative video. Points are color-coded according to each estimate’s likelihood. Only grayscale points (LH > .75) were “confirmed” and used for classification. No footfalls with this paw were classified as errors. **C**. As in panel B, for the back left paw. Two footfall errors were detected, filled in green. Errors began with the first confirmed position before threshold, and ended with the first confirmed position above baseline. **D.** Estimated interpolated velocity for confirmed foot positions. Also shown is a representative curve (generalized additive model, solid gray line), and overall mean velocity (total distance/total time, dashed green line). **E.** Footfall errors (circles = females, diamonds = males) per week by the control (pink) and experimental (blue) groups. Bootstrapped means are reported (bars = 80% confidence intervals, whiskers = 95% confidence intervals). Points represent discrete counts, but were randomly jittered on both axes to assist visual inspection. **F.** Summary of estimated error rates for an average subject in each condition (green = baseline, pink = control, blue = experimental), for both sexes (darker colors = female, lighter colors = male). Posterior estimates were computed using Equations 1 and 2, limited to only fixed effects (all *β* terms) and the population mean across subjects (*η*). Estimates for the control and experimental conditions here averaged performance across all six weeks (bars = 80% posterior credible intervals, whiskers = 95% credible intervals). **G.** As in panel E, but plotting overall velocities (pixels/second). Points were jittered on the horizontal axis to assist visual inspection. **H.** As in panel F, but plotting overall velocities (pixels/second).

**S3 Fig 3. Week-by-week estimates of performance. A.** Estimated error rate for an average subject during each week, split by condition (green = baseline, pink = control, blue = experimental) and sex (darker colors = female, lighter colors = male), based on the population-level effects in Equations 1 and 2 (bars = 80% posterior credible interval, whiskers = 95%). **B.** Posterior mean estimates of individual error rates per session for each female subject, based on the full model in Equations 1 and 2. Each point reflects one subject during that session. Subjects are only plotted for weeks where their data are available. Trendlines show the overall linear trend across subjects for the weeks after baseline (shaded region = 95% credible interval). **C.** As in panel C, but plotting male subjects. **D-F.** As in panels A-C, but plotting estimated velocities (pixels/second).

**S4. Sample Video – DLC Tags.** Example video showing nine body points labeled over time during a single recording session.

**S5. Sample DLC Raw Data.** Raw data extracted from a single session video, including the x and y coordinates and the estimated likelihood for each body tag across frames.

